# Screening channelrhodopsins using robotic intracellular electrophysiology and single cell sequencing

**DOI:** 10.1101/2025.08.19.671087

**Authors:** Samuel Ehrlich, Alexandra D VandeLoo, Mohamed Badawy, Mercedes M. Gonzalez, Max Stockslager, Aimei Yang, Sapna Sinha, Shahar Bracha, Demian Park, Benjamin Magondu, Bo Yang, Edward S. Boyden, Craig R. Forest

## Abstract

**Background:** Our ability to engineer opsins is limited by an incomplete understanding of how sequence variations influence function. The vastness of opsin sequence space makes systematic exploration difficult.

**New method:** In recognition of the need for datasets linking opsin genetic sequence to function, we pursued a novel method for screening channel-rhodopsins to obtain these datasets. In this method, we integrate advances in robotic intracellular electrophysiology (<u>P</u>atch) to measure optogenetic properties (<u>E</u>xcite), harvest individual cells of interest (<u>P</u>ick) and subsequently sequence them (<u>S</u>equence), thus tying sequence to function.

**Results:** We used this method to sequence more than 50 cells with associated functional characterization. We further demonstrate the utility of this method with experiments on heterogeneous populations of known opsins and single point mutations of a known opsin. Of these point mutations, we found C160W ablates ChrimsonR’s response to light.

**Conclusion and comparison to existing methods:** Compared to traditional manual patch clamp screening, which is labor-intensive and low-throughput, this approach enables more efficient, standardized, and scalable characterization of large opsin libraries. This method can enable opsin engineering with large datasets to increase our understanding of opsin sequence-function relationships.

## 1. Introduction

Opsins are a family of light-sensitive proteins that enable precise control of cellular activity and have become indispensable tools for neuroscience, transforming our ability to dissect and manipulate neural circuits (1; 2). Their utility depends on the inherent properties of each specific channel (such as photocurrent amplitude, spectral response, and channel kinetics (3)), which are in turn determined by the protein’s amino acid sequence. Engineering opsins with improved or novel properties is essential for improving their utility in both basic science and clinical settings. Improved properties have been achieved by rational design (4, 5, 6, 7, 8), exploring the natural diversity of opsins (9, 10), machine learning (11, 12), and directed evolution (13).

Although in the last two decades many optogenetic tools have been developed there are still many open frontiers and unmet needs. For example, opsins with enhanced sensitivity for low-light activation (to reduce toxicity and tissue heating), spectrally-shifted for deeper tissue penetration and less cross-talk with other tools, reduced immunogenicity for long-term expression and more. Our ability to engineer opsins to meet these needs is limited by an incomplete understanding of how sequence variations influence function and the vastness of the genetic sequence space makes systematic exploration difficult with current tools like patch clamp electrophysiology.

Patch clamp recording is the gold standard method for resolving millivolt- and millisecond-scale dynamics and ion conductances of light activated channels (14). Consequently, laborious opsin screens that incorporate patch clamp electrophysiology have produced rich data sets to drive further opsin discovery (9). Prior work has improved the throughput and efficiency of patch clamping through new techniques such as pipette cleaning (15) and automation (16, 17, 18, 19).

Single-cell sequencing technologies have dramatically expanded our ability to link genotype with phenotype at cellular resolution. In neuroscience and molecular biology, these methods have enabled the reconstruction of transcriptional profiles, identification of rare cell types, and resolution of heterogeneous responses within genetically similar populations (20, 21, 22). When coupled with functional assays, single-cell sequencing allows direct attribution of observed cellular behavior—such as ion channel kinetics or optogenetic response profiles—to the underlying gene sequence. This capability is particularly impactful for protein engineering, where high-throughput sequencing can validate the identity of functionally characterized cells and support genotype-phenotype mapping.

Recognizing that there is a need for datasets linking opsin genetic sequence to function, we pursued a novel method for screening opsins to obtain these datasets. In this method, we integrate advances in robotic intracellular electrophysiology (<u>P</u>atch) to measure optogenetic properties (<u>E</u>xcite), harvest individual cells of interest (<u>P</u>ick) (23, 24) and subsequently sequence them (<u>S</u>equence), thus tying sequence to function. The PEPS method enables direct, high-resolution screening of opsin libraries transfected into mammalian cells. We demonstrate the capabilities of this system through a series of experiments on a homogeneous population of ChrimsonR expressing cells, a heterogeneous population of known opsins (ChrimsonR and CheRiff), and a population of unknown ChrimsonR mutants. This system can enable opsin engineering with large datasets to increase our understanding of opsin sequence-function relationships.

## 2. Materials and methods

### 2.1. Cell culture

HEK293FT cells (Invitrogen) were selected as the expression host for several reasons: (i) they are highly amenable to calcium phosphate and lipofectamine transfection; (ii) they are widely used for robust expression of exogenous proteins, making them ideal for optogenetic tool characterization; (iii) they are reported to have among the lowest rates of exogenous DNA-induced mutation among commonly used mammalian cell lines; (iv) they are easy to grow and maintain, and widely adopted for tool development and screening.

Cells were cultured at 37°C with 5% CO_2_ in DMEM (Gibco) supplemented with 10% heat-inactivated fetal bovine serum (FBS, Corning), and 1% sodium pyruvate (Gibco). Cells were maintained between 10% and 70% confluence to preserve exponential growth. Cells were seeded onto coverslips (0.15 mm thick, 12 mm in diameter, Electron Microscopy Systems) coated in Matrigel (2% growth factor–reduced Matrigel, Corning) diluted in DMEM for 1 h at 37 °C.

For the heterogeneous experiment (Section 3.3) cells transfected with either ChrimsonR or CheRiff separately were trypsinized from coverslips on the day of the experiment, mixed in equal proportions, then seeded to new coverslips. For the library experiment described in Section 3.4, cells were trypsinized and mixed in a similar fashion as Section 3.3. This procedure ensured co-culture of cells expressing different opsin constructs on the same recording surface and was identical in preparation to the heterogenous experiment.

### 2.2. Cell transfection

We used the pN1 expression vector (Clontech), which contains an SV40 origin of replication, allowing episomal replication in HEK293FT cells. This enhances protein expression even under single-copy plasmid delivery. The CMV promoter was used to drive expression of all constructs due to its established high strength in HEK293-derived lines. For experiments described in Section 3.2 and Section 3.3, 10,000 cells were seeded onto the coverslips and transfections were performed the following day using a commercially available calcium phosphate transfection kit (Invitrogen). We combined 150 ng of DNA, 3.125 µl of CaCl_2_, 25 µl of HBS, and water for a total volume 50 µl and incubated for 20 min. 25 µl of solution was then added to each individual well. After two days of incubation the experiment was performed. For the heterogeneous experiment (Section 3.3), cells were incubated with either ChrimsonR or CheRiff separately, trypsinized from coverslips on the day of the experiment, mixed in equal proportions, then seeded to new coverslips. For experiments described in Section 3.4, approximately 70,000 cells were seeded onto the coverslips and transfected the following day using 1 µl of Lipofectamine 2000 (Invitrogen) and 150 ng of DNA. After one day of incubation the cells were trypsinized and placed on new coverslips at a lower density. Mutant plasmids were ordered from Azenta Life Sciences.

### 2.3. Microscope

All experiments were performed using an inverted fluorescence microscope (Nikon Eclipse Ti2000) configured for simultaneous patch clamp recording, light stimulation, and single-cell harvesting. A 40× objective (Nikon, Plan Fluor, NA 0.75) was used for both fluorescence imaging and whole-cell patch clamping. Fluorescence images were captured with a pco.panda 4.2M scientific CMOS camera (PCO AG, Germany).

The microscope was mounted on a motorized XY stage (Scientifica) for precise positional control. Z-axis focus control was achieved via a stepper motor (Scienctifica) coupled to a high-precision gear box radially coupled to the objective focus wheel, allowing automated adjustments of focal plane during pipette approach and cell imaging.

### 2.4. Electrophysiology instrumentation

All patch clamp recordings were conducted using an Axopatch 200B amplifier (Molecular Devices). The analog voltage output from the amplifier was digitized and recorded using a National Instruments data acquisition device (NI USB-6210, National Instruments), which provided synchronized analog-to-digital conversion and timing control for stimulus delivery and data acquisition.

The patch pipette was mounted on a three-axis micromanipulator (Sensapex), which enabled fine automated positioning and alignment for seal formation and whole-cell configuration. Fluorescence excitation for identifying opsin-expressing cells was provided by a mercury arc lamp (X-Cite 120, Excelitas Technologies), with standard filter sets for tdTomato and eGFP (Thermo Fischer Scientific).

Optogenetic stimulation was delivered through the patch pipette using an LED light engine (Lumencor Spectra X), which provided illumination across multiple wavelengths. Light was coupled into the back of the patch pipette through an optical fiber inserted into a special purpose pipette holder (Optopatcher) (25), enabling precisely colocalized stimulation and recording. Pressure control for the patch pipette was maintained using a custom-built multi-channel pressure control box that delivered computer-regulated positive and negative pressures for cell access and seal formation as previously described (17). A single pipette was reused unless it failed due to clogging, breaking, or repeated failure to form a gigaseal.

### 2.5. Electrophysiology recording

The external (bath) solution was Tyrode’s solution, containing (in mM): 125 NaCl, 2 KCl, 3 CaCl_2_, 1 MgCl_2_, 10 HEPES, and 30 glucose (pH adjusted to 7.3 with NaOH, 305 mOsm). The internal pipette solution contained (in mM): 125 K-gluconate, 8 NaCl, 0.1 CaCl_2_, 0.6 MgCl_2_, 1 EGTA, 10 HEPES, 4 Mg-ATP, and 0.4 Na-GTP (pH adjusted to 7.3 with KOH, 295–300 mOsm). For recording, we manually selected cells exhibiting the following criteria:

1. non-zero fluorescence visible at set exposure (1 sec on our camera) and
2. physically isolated from neighboring cells. Whole-cell patch-clamp recordings were performed on isolated HEK293FT cells to minimize space-clamp artifacts and ensure accurate measurements of photocurrents. All recordings were conducted at room temperature. All data was collected with a 10 kHz lowpass Bessel filter applied.

Patched cells which did not meet the following patch quality control thresh holds were rejected from analysis: access resistance <35 MΩ and holding current within *±*100 pA. Typical membrane resistance ranged from 200 MΩ to 2 GΩ, and pipette resistance was 3–9 MΩ. Patch pipettes were pulled from borosilicate glass with a 1.5 mm outside diameter and a 0.86 mm inside diameter (Warner Instruments).

Each cell underwent a brief voltage-clamp test step of +10 mV from holding potential to assess membrane capacitance and seal quality. This was followed by a current-clamp protocol consisting of three consecutive current injections (−10, 0, and +10 pA) to confirm membrane responsiveness and resting potential stability. After patching, pipettes were cleaned as previously described using tergazyme (Alconox) (15).

#### 2.5.1. Optical stimulation

To determine the optical stimulation protocol we first found the minimum illumination power needed to produce the maximum photocurrent amplitude of the ChrimsonR sequence and introduce a safety factor in order to reduce the risk of photobleaching during experiments. This calibration is done by performing intensity sweeps (A.6) to determine the optimal illumination power at the 635nm wavelength. We chose to set the illumination power at approximately 75% of the power at maximum sustained photocurrent amplitude for the experiments herein (0.842 mW as measured at the end of the optical fiber). The other excitation wavelengths were calibrated to match the photon flux of the chosen power for the 635 nm light.

The light protocol consisted of three randomized sweeps per cell, each containing six discrete wavelengths: 395, 440, 470, 575, 635, and 740 nm. For each sweep, cells were held in darkness for 15 sec, stimulated for 1 second with a single wavelength, and returned to darkness for 14 sec, totaling 90 sec per full stimulation series while being held by voltage clamp at −50 mV. Wavelengths were randomized across trials to control for adaptation and order effects. All light intensities were pre-calibrated for consistent photon flux delivery across wavelengths.

### 2.6. Cell harvesting

Following patch-clamp electrophysiology recordings, individual cells were harvested for sequencing using the CellSorter system (CellSorter Inc.) mounted vertically above the microscope stage. The CellSorter combines a vertical single-axis micromanipulator and independent pressure controller for precise cell aspiration and deposition.

For collection, pulled glass micropipettes with an inner diameter of 70 µm, a tip length of 5 mm, and a pull length of 80 mm were used. The target cell was approached until the tip of the pipette was 15 µm above the cell as determined by calibration at the cell focal plane. Negative pressure was valve actuated for 100 ms by drawing a 60 ml syringe to 40 ml, producing a brief and controlled suction event and aspirating the cell into the pipette tip. The collected cell was deposited into a PCR tube containing 10 µl of QuickExtract DNA Extraction Solution (Lucigen). To expel the cell and associated buffer, 10 µl of liquid was discharged from the pipette using positive pressure. Positive pressure was delivered by a gravity-fed water reservoir (40 mm Hg) connected to the back of the pipette holder via tubing, allowing valve actuated release of pressure.

### 2.7. Software

We designed and built a custom software suite to operate the integrated robotics system used for automated patch clamp recording and single-cell harvesting based on previous work (17). The software coordinates all aspects of the experimental workflow, including pipette positioning, pressure control, light stimulation, electrophysiological recording, and post-recording cell collection.

The software interfaces with all hardware components through vendor-provided and custom-built drivers. Each component is synchronized through a unified interface, allowing precise control of experimental timing and coordination.

LabVIEW (National Instruments) was used as the primary software environment for hardware integration and experimental control. It was selected due to its robust support for real-time data acquisition, digital and analog I/O, and synchronization across multiple hardware platforms. All manipulator motion, light triggering, pressure switching, and amplifier communications were controlled via LabVIEW, which also hosted the central graphical user interface used during experiments.

MATLAB (MathWorks) was used for real-time data analysis and feedback. During patch clamp recordings, MATLAB scripts monitored membrane and access resistance, holding current, and seal quality in real time. This information was relayed back to the LabVIEW controller to support automated decision-making for gigaseal formation, break-in, and whether to proceed with optical stimulation and cell harvesting.

The integrated software supports real-time monitoring of key electrophysiological parameters, enabling automated decision-making for seal success and break-in. Following successful recording, the cell is collected via a second robotic manipulator (CellSorter API) and deposited into individual collection tubes for downstream sequencing. All experimental parameters, including manipulator motion, light delivery, pressure regulation, and data acquisition, are synchronized through the central software interface. This system enables semi-autonomous operation of the patch-and-harvest pipeline, significantly reducing operator input.

### 2.8. Single cell sequencing

To lyse the collected cells and release plasmid DNA for downstream amplification and sequencing, the single-cell containing collection tubes were processed using a PCR-compatible thermal lysis protocol. Each tube was vortexed for 30 sec, then placed in a thermocycler and incubated at 65°C for 6 min. Following incubation, the samples were vortexed again for 30 sec, then heated at 98°C for 2 min to complete the lysis.

After thermal lysis, 2.5*µ*l of TE buffer (10mM Tris-HCl, 1mM EDTA, pH 8.0) was added to each tube to dilute and stabilize the lysate for downstream processing. Lysates were stored at –20°C until further use.

A 7*µ*l aliquot of the lysate was used as the template for standard PCR, using the following primers: Forward (5): GCAGAGCTGGTTTAGTGAACC and Reverse (3): CCGCATGAACTCTTTGATCACTTCC. Then 10*µ*l portion of the resulting PCR product was submitted to Plasmidsaurus for Nanopore sequencing.

### 2.9. Data analysis

Electrophysiology data was analyzed using custom made Python scripts. All data was filtered with a digital 3 kHz Bessel filter after data collection. Sustained photocurrents were defined as the average of the photocurrent from the baseline holding current during the last 50 ms of light stimulation. For experiments with known opsins in homogeneous or heterogeneous populations, only cells which exhibited sustained photocurrents greater than 15 pA were included in spectral response analysis. Fluorescent expression was determined using ImageJ. Genetic sequences were aligned using the Smith-Waterman method in SnapGene. Error bars indicate the standard error of the mean (SEM) in all figures.

## 3. Results

### 3.1. System overview

To enable direct, sequence-resolved functional screening of opsin libraries, we designed and implemented the Patch, Excite, Pick, Sequence (PEPS) method (Fig. 1). The PEPS method encompasses a cycle of steps through which optogenetic function and sequence datasets are obtained (See Supplementary Video). Following functional characterization, each individual cell is immediately harvested directly from the coverslip using a second robotic manipulator equipped with a dedicated harvest pipette. The harvested cell is deposited into a collection tube for downstream single-cell sequencing, linking each cell’s optogenetic response profile to its underlying plasmid sequence. Once a maximum of 10 cells have been harvested, the cells are removed and put through a thermocycler protocol to prepare for subsequent sequencing. This approach allows for characterization of opsin variants at the single-cell level and direct recovery of individual cells for sequencing, supporting genotype-phenotype mapping.

**Figure 1:**
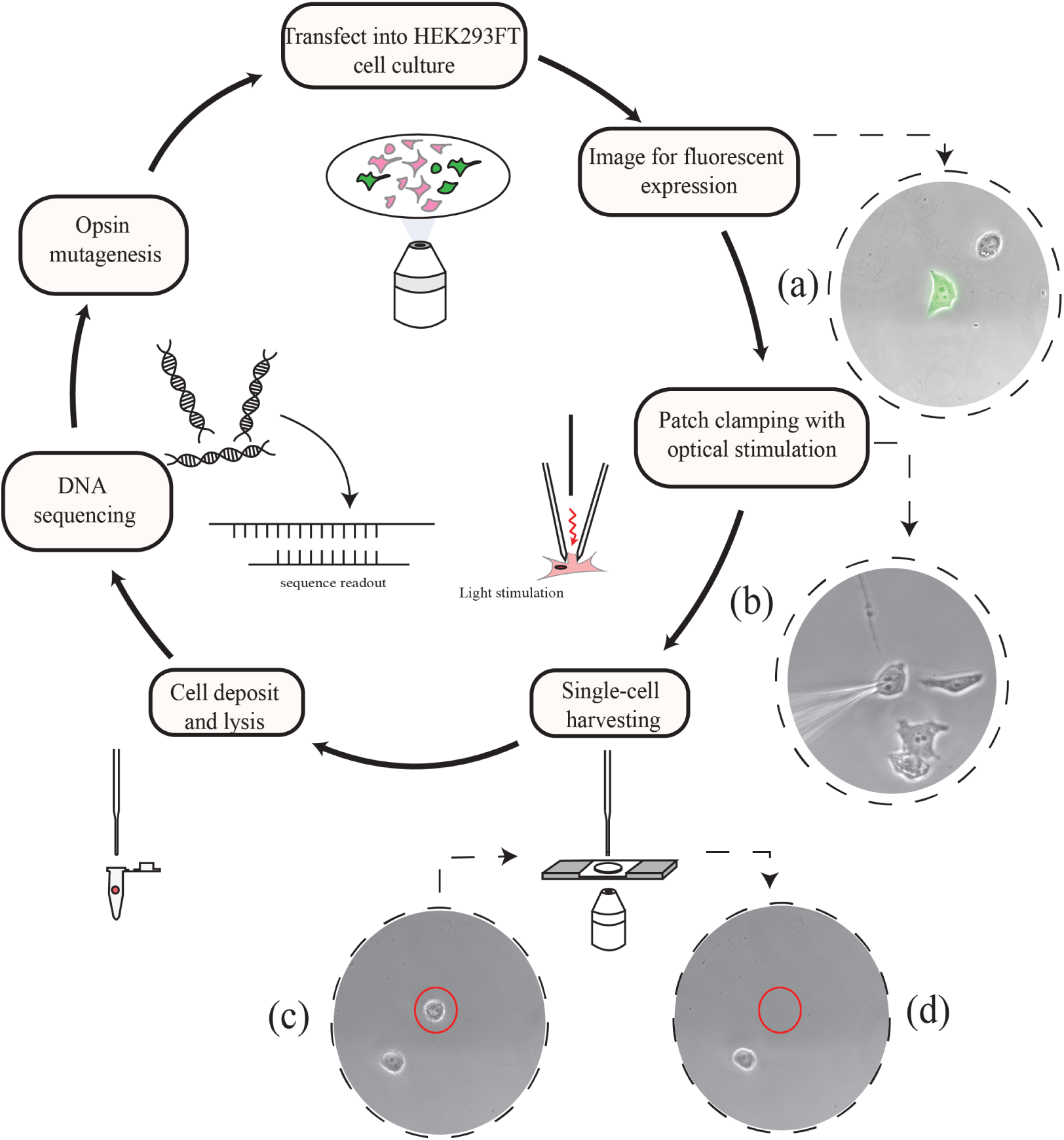
The patch, excite, pick and sequence (PEPS) pipeline. An opsin plasmid is transfected into HEK293FT cells. After 1-2 days, fluorescence images reveal cell expression. Cells that are isolated and fluorescent (see photograph in inset (a)) are optically stimulated while intracellular electrophysiology is recorded (see photograph in inset (b)). Successfully patched cells are harvested with suction from a pipette that cleanly removes the cell from the coverslip (see photographs in insets (c) and (d)) Each cell is deposited and lysed followed by sequencing.

The instrument for steps from image fluorescent expression to cell deposit and lysis is shown in Fig. 2. Key subsystems corresponding to steps in Fig. 1 are shown in Fig. 2A, with details of the embodiment in Fig 2B.

**Figure 2:**
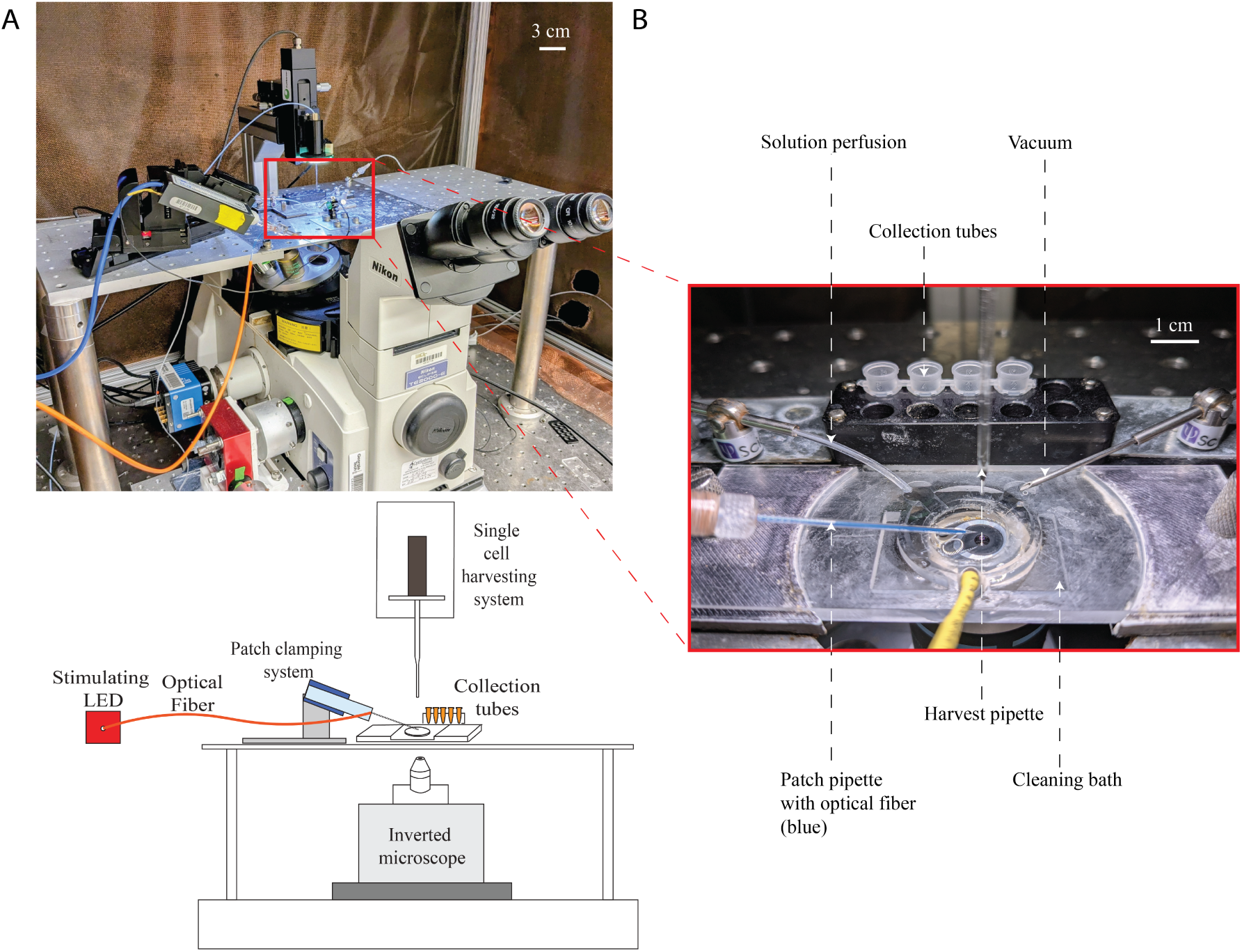
Overview of the PEPS hardware (A) Photograph and schematic (for clarity) illustrating the key subsystems of the robot that enables patch clamp electrophysiology, optogenetic stimulation via the stimulation LED and optical fiber, and cell harvesting system all above an inverted fluorescence microscope for imaging. (B) Closeup photograph showing interaction and orientation of tubing for bath perfusion and aspiration, pipettes used for patching and harvesting, and pipette cleaning bath for pipette reuse.

### 3.2. Homogeneous population of known opsin-ChrimsonR

To assess the overall efficiency of the PEPS method, we tracked the outcome of 176 patching attempts targeting ChrimsonR-expressing cells (Fig. 3A). Of these, 137 resulted in gigaseal formation and 119 proceeded to break in. Cells were further filtered for quality based on access resistance and holding current, resulting in 100 quality whole cells after 5 whole cells were lost during the photostimulation. Among these, 63 cells were successfully picked up using the automated CellSorter module, and 45 were ultimately sequenced (for reference sequence see Fig. A.8). This corresponds to an overall success rate of 26% from patch attempt to sequence data, with each stage of the process contributing a quantifiable efficiency.

**Figure 3:**
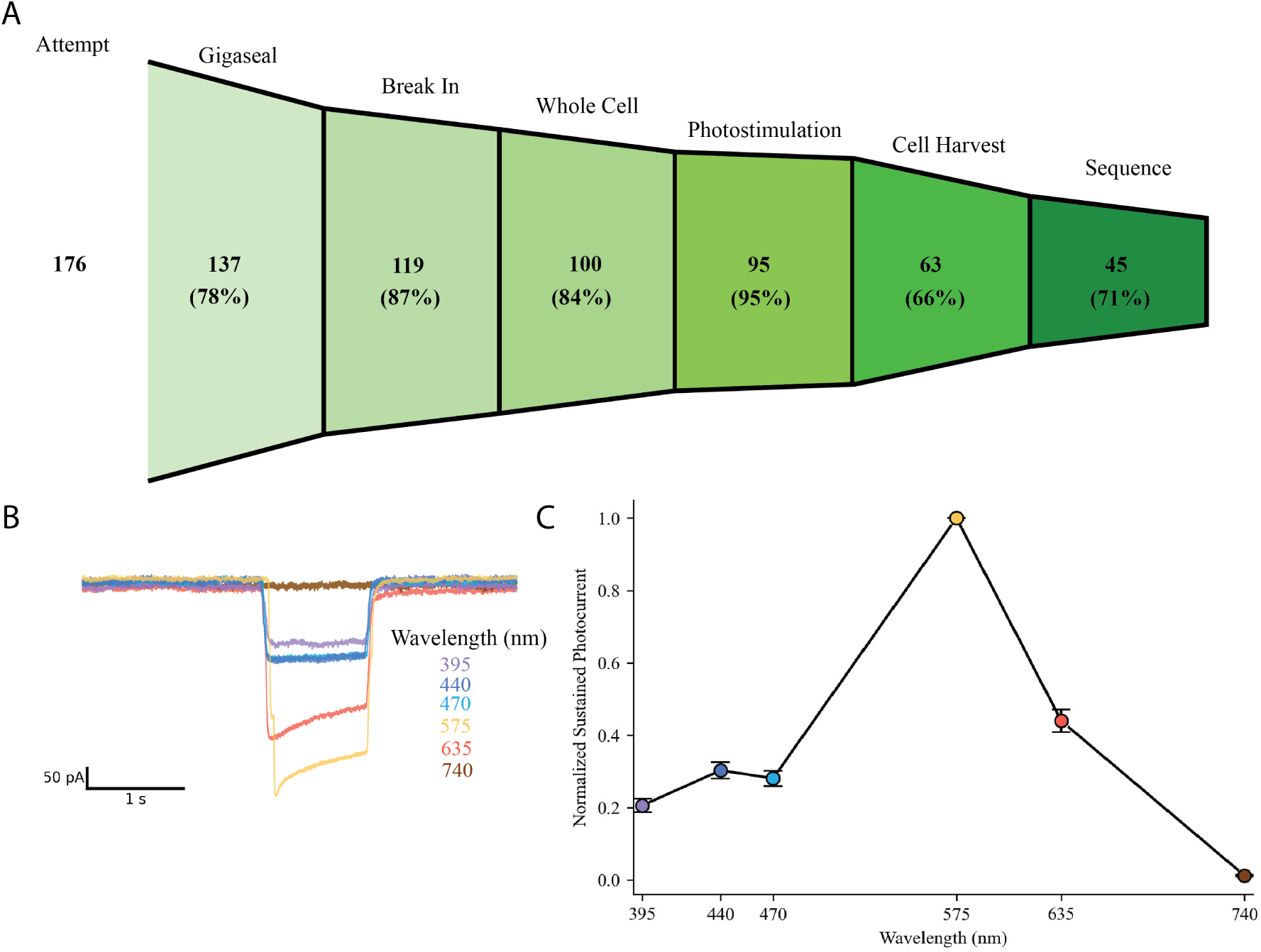
Electrophysiological characterization and throughput metrics for ChrimsonRtdTomato expressing cells. (A) Funnel diagram illustrating experimental efficiency through each stage of the PEPS workflow. Of 176 total patching attempts, 137 resulted in gigaseals, 119 reached whole-cell configuration, 100 met quality control criteria, 95 maintained the patch through photostimulation, 63 were harvested, and 45 were successfully sequenced. (B) Representative whole-cell patch clamp recordings of ChrimsonR-tdTomato-expressing HEK293FT cell stimulated at six different wavelengths (395, 440, 470, 575, 635, and 740 nm) using matched photon flux. (C) Normalized sustained photocurrent responses across wavelengths reveal peak activation at 575 nm, consistent with ChrimsonR’s known spectral sensitivity. Error bars represent SEM (n = 22).

Each successful patch and subsequent harvest takes about 18 min (up to 2 min to gigaseal, 1 min to break in, 1 min to stabilize, 10 min for electrophysiology measurements, 2 min to clean the pipette, 2 min for collection and deposition). Including time to find an appropriate cell to patch, this gives a theoretical maximum of 3 cells per hour for this protocol. It should be noted that 50% of the duration is due to the stimulation protocol used in this work and could vary significantly depending on application.

To evaluate our ability to screen channelrhodopsins using the PEPS method, we first conducted a survey of HEK293FT cells expressing ChrimsonR-tdTomato. Representative whole-cell patch clamp recordings revealed photocurrent responses from ChrimsonR-expressing cells across the range of stimulation wavelengths (Fig. 3B) with 575 nm producing the maximum sustained photocurrent with magnitudes as large as 350 pA (Fig. 3C). Sequence data for all 45 cells from the homogenous population are identical to the example sequence shown in Fig. A.8.

### 3.3. Heterogeneous population of known opsins-ChrimsonR and CheRiff

To assess the capability of the PEPS method to distinguish opsin variants within a heterogeneous population, we prepared a cell population expressing either ChrimsonR-eGFP or CheRiff-eGFP. First, control data was collected on homogeneous populations of both ChrimsonR (n=11) and CheRiff (n=11) to define the expected response. Next, cells were transfected with ChrimsonR or CheRiff separately, then trypsinized and mixed together to form a heterogeneous population of known opsins. Critically, the opsin being expressed by a particular cell could not be identified during the experiment by fluorescence imaging alone since the fluorescence tag is identical for these two constructs. Whole-cell voltage-clamp recordings were acquired from selected fluorescent cells and analyzed for spectral response (Fig. 4A). The distinct spectral signatures enabled classification of each cell based solely on this functional data. The spectral profiles segregate into two clusters consistent with ChrimsonR and CheRiff reference data (Fig. 4B). Sequencing data from each picked cell in the heterogeneous population (n=6) independently confirmed expression of either ChrimsonR or CheRiff sequences (See Fig. A.8 and A.9 for example sequences). These results validate the use of the PEPS method to distinguish between opsins by spectral properties as well as by sequence data. Efficiency and throughput were similar to the previous experiment (not shown).

**Figure 4:**
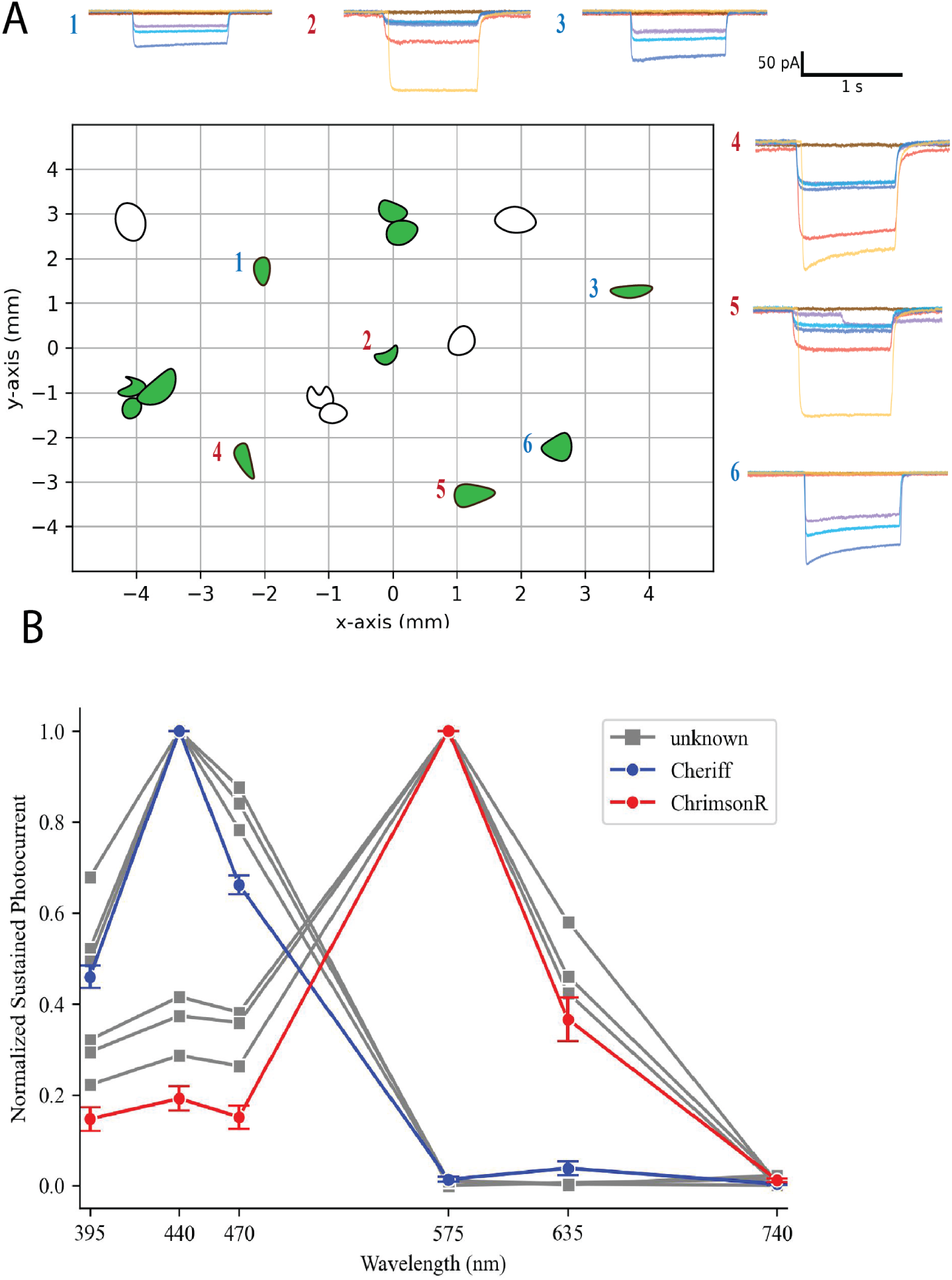
Identification of opsins from spectral signatures in a hetergenous population (A) Spatial map of patched cells overlaid with representative photocurrent traces. Cells 2, 4, and 5 display ChrimsonR-like responses (red numbers), while cells 1, 3, and 6 exhibit CheRiff-like responses (blue numbers). (B) Normalized sustained photocurrent responses across wavelengths reveal peak activation at 575 nm for ChrimsonR and 440 nm for CheRiff, with the unknown sequences from the heteregenous population clustering with either ChrimsonR or CheRiff. Error bars represent SEM (n = 11 for ChrimsonR, n=11 for CheRiff, n=6 for unknown). 15

### 3.4. Heterogeneous population of unknown opsins-ChrimsonR mutant library

To test the ability of our PEPS method to resolve functional properties associated with individual mutations, as would be necessary to generate a function-sequence dataset, we constructed a small opsin library containing single amino acid substitutions to ChrimsonR. The selected mutations (C160W, E300L, F341E, H291Y, and H307L) span distinct regions of the channel pore and transmembrane helices, illustrating the potential diversity of mutation sites within a larger, unbiased mutant library (Fig. 5A). Residue E300 is a known important site in the ChrimsonR ion gate (26); F341 would not be expected to change function as it lies on the terminal end of the protein in a disordered region, and H291 is a conserved residue across several opsins (26) and would be expected to impact function. C160 and H307 are random locations. This approach mimics a realistic mutagenesis strategy for directed evolution, in which single or combinatorial point mutations are introduced across the coding sequence to probe sequence–function relationships.

**Figure 5:**
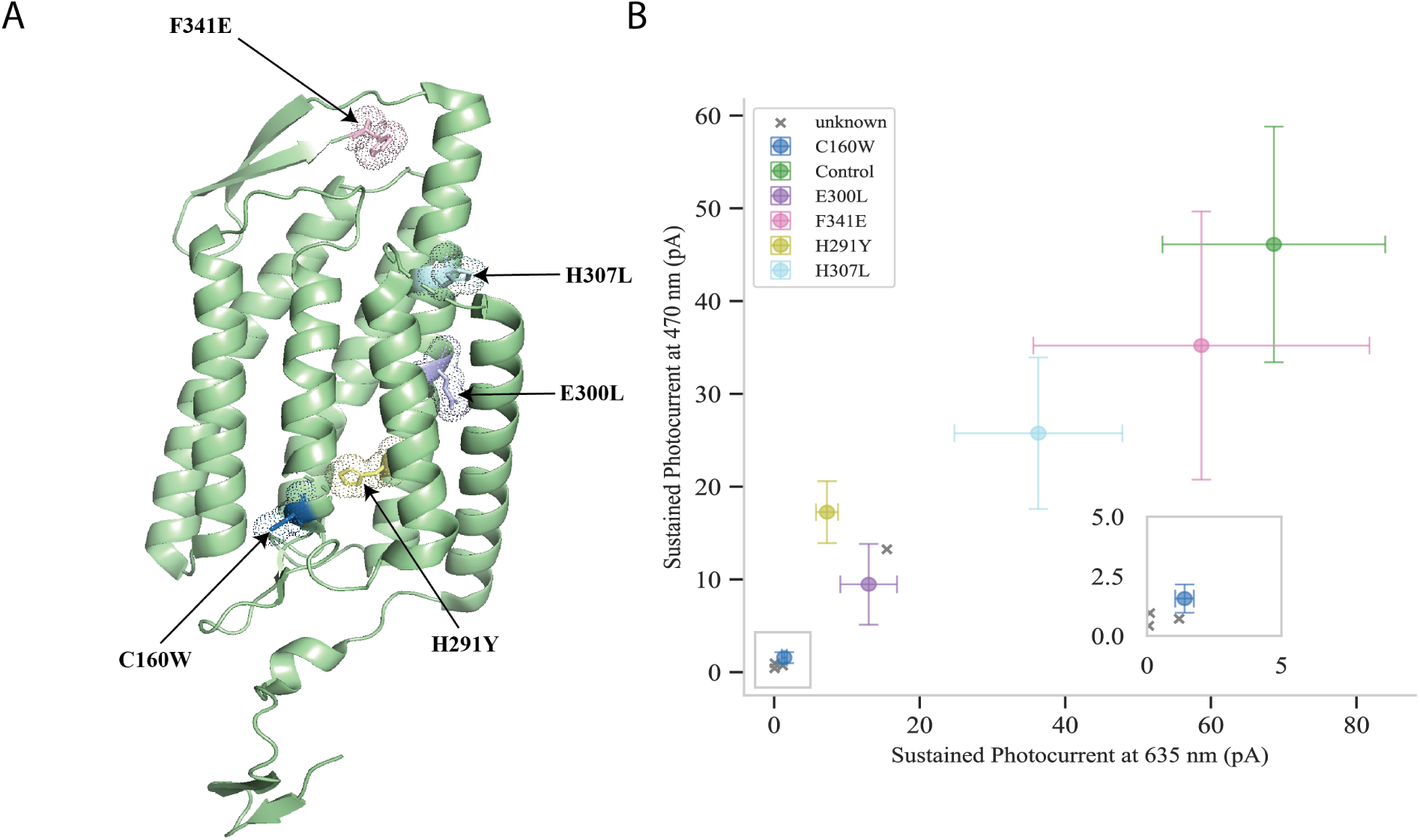
Single amino acid substitutions and functional mapping of ChrimsonR variants. (A) Crystal structure model of ChrimsonR highlighting the positions of five single amino acid substitutions (C160W, E300L, F341E, H291Y, H307L) introduced to generate a representative mutant library. (B) Scatter plot showing the mean sustained photocurrent amplitudes at 470 nm versus 635 nm for control cells (labeled) and unknown cells harvested from a mixed population. Error bars represent SEM. Unknown cells were sequenced and matched to the indicated single mutants, demonstrating the system’s ability to resolve functional variation and associate phenotype with genotype. (n = 6 for control populations, n=4 for unknown.

To establish control photocurrent responses for each variant in a homogeneous population, we measured the sustained photocurrent across the six wavelengths (395–740 nm) for each single mutant in HEK cells (Fig. 5B; Appendix A.10). Note that Fig. 4B represents the spectral response data differently than in the previous experiments to highlight the change in photocurrent between mutants, since the full spectra are more difficult to interpret when the change is relatively small. As expected, the different single substitutions produced a range of photocurrent amplitudes and spectral responses. Notably, C160W ablated photocurrents across all wavelengths relative to other variants and the parental ChrimsonR.

We then mixed a population of cells containing these mutants, recorded their photocurrent responses, and harvested individual cells for genotyping.

Four cells made it through the entire pipeline and are represented in Fig. 5B as “unknown”. Upon sequencing these unknown cells, three expressed the C160W mutation while one expressed the E300L mutation (See Appendix A.11 and A.12). The photocurrents of the sequenced cells showed close alignment with the control populations (Fig. 5B). The distribution of these unknown cells within the scatter plot highlights the variability inherent in patch clamp photocurrent recordings, which can be influenced by cell health, expression level, and recording conditions, among others.

## 4. Discussion

We present a method that combines robotic intracellular patch-clamp electrophysiology with single-cell harvesting and sequencing to link opsin genetic sequence to functional performance that we term Patch, Excite, Pick, Sequence (PEPS). By integrating and automating each step—from whole cell patch clamping and recording to harvesting, this system addresses key limitations in traditional opsin screening workflows. This includes the need for labor-intensive patching and the difficulty of connecting precise functional measurements to sequence information in a scalable manner. We first demonstrated PEPS on a homogenous population of cells expressing ChrimsonR and measured the throughput of the method. Of attempted cells, 26% made it through the entire process. The spectral response of the ChrimsonR expressing cells matched previous findings with 575 nm light producing the maximum sustained photocurrent with our system (9, 26).

We observed considerable variability in photocurrent amplitudes among ChrimsonR-expressing cells, even under uniform stimulation. The sustained photocurrent measured did not correlate with the level of fluorescent expression (Fig. A.7) as has been found in other work (9). Cells with low or undetectable fluorescence exhibited negligible or absent photocurrent, suggesting that a minimum expression level is a necessary but not sufficient determinant of functional response. These data support the inclusion of minimum fluorescence thresholds when selecting cells for patching in future library screens.

After testing PEPS with a homogenous population, we next tested a heteregenous population of cells expressing either ChrimsonR or CheRiff. Cells showed a distinct spectral response and subsequent sequencing confirmed either the presence of ChrimsonR or CheRiff. This demonstrated that PEPS could be used on a hetergenous population to functionally characterize individual cells and tie that functional data to the sequence.

In our final experiment, we tested a small sample library of single amino acid substitutions in ChrimsonR to mimic conditions that would be expected in a protein engineering campaign. The sustained photocurrents of the cells that went through the entire method aligned closely with the homogeneous control population of their respective mutant.

Importantly, while absolute photocurrent amplitudes are inherently variable in this pipeline, this variability does not preclude meaningful interpretation on a qualitative level. For example, none of the sequenced C160W cells produced photocurrents matching the levels observed for the functional wild type control, reinforcing that this mutation likely results in nonfunctional channels. This binary readout—functional versus nonfunctional—has previously proven valuable in informing machine learning models for protein engineering, where clear classification of variant function can provide robust training data (12). Together, these results demonstrate that our system can link electrophysiological measurements to single-cell genotypes, enabling direct assessment of functional consequences for individual amino acid substitutions in opsins.

Related approaches, such as Patch-seq, have successfully combined intracellular recordings with single-cell transcriptomics to map cell types and relate gene expression profiles to electrophysiological properties. Patch-seq has been instrumental in neuroscience for uncovering cellular diversity within the brain by recovering mRNA transcripts after patching (27). In contrast, our method leverages the fact that the target gene is encoded on plasmid DNA within the nucleus of transfected HEK cells. By harvesting the entire cell—including its genomic and plasmid DNA—we avoid the technical challenges and potential biases introduced by RNA extraction and reverse transcription. This change reduces contamination risks, facilitates automation, and enables robust genotyping of expressed opsin constructs. While Patch-seq remains valuable for cell type classification and endogenous transcriptome analysis, especially for cells embedded in tissue, our approach is purpose-built for protein engineering applications where the priority is to recover the sequence of the target variant.

Looking forward, the method described here can be scaled and adapted for broader applications in opsin engineering and beyond. The system provides a foundation for opsin engineering campaigns that require reliable, high-resolution functional readouts that cannot be captured with optical voltage indicators alone. Future extensions could integrate more complex stimulation protocols to probe additional properties such as channel kinetics, recovery from inactivation, or ion selectivity. Likewise, this approach could be adapted to other classes of membrane proteins where patch clamp measurements are essential but library screening remains impractically low-throughput with traditional workflows.

The ability to directly link sequence and function at the single-cell level will be particularly valuable for training and validating machine learning models that aim to navigate vast protein sequence landscapes. By combining unbiased mutagenesis, robust electrophysiological measurement, and single-cell sequencing, our platform can generate comprehensive datasets that enhance our understanding of opsin sequence–function relationships and accelerate the design of new optogenetic tools with tailored properties for neuroscience and beyond.

## Acknowledgments

CRF acknowledges the NIH BRAIN Initiative Grant (NEI and NIMH 1-U01-MH106027-01), NIH R01NS102727, NIH Single Cell Grant 1 R01 EY023173, NIH R01DA029639 and NIH RF1AG079269, support from Georgia Tech through the Institute for Bioengineering and Biosciences, Invention Studio, and the George W. Woodruff School of Mechanical Engineering. S.S. acknowledges the Schmidt Science Fellows for their generous support through the postdoctoral fellowship.

## 5. Author contributions

**SE**: Conceptualization, Data curation, Formal analysis, Investigation, Methodology, Software, Validation, Visualization, Writing – original draft, Writing – review and editing. **ADV**: Conceptualization, Writing – original draft, Writing – review and editing. **MB**: Investigation, Methodology, Software. **MG**: Investigation. **MS**: Methodology, Software. **AY**: Methodology. **SS**: Writing – review and editing. **SB**: Writing – review and editing. **DP**: Methodology. **BM**: Methodology. **BY**: Methodology. **ESB**: Conceptualization, Funding acquisition, Methodology, Supervision, Writing – review and editing. **CRF**: Conceptualization, Data curation, Formal analysis, Funding acquisition, Methodology, Supervision, Writing – original draft, Writing – review and editing.

## Appendix A. Supplementary material

**Figure A.6:**
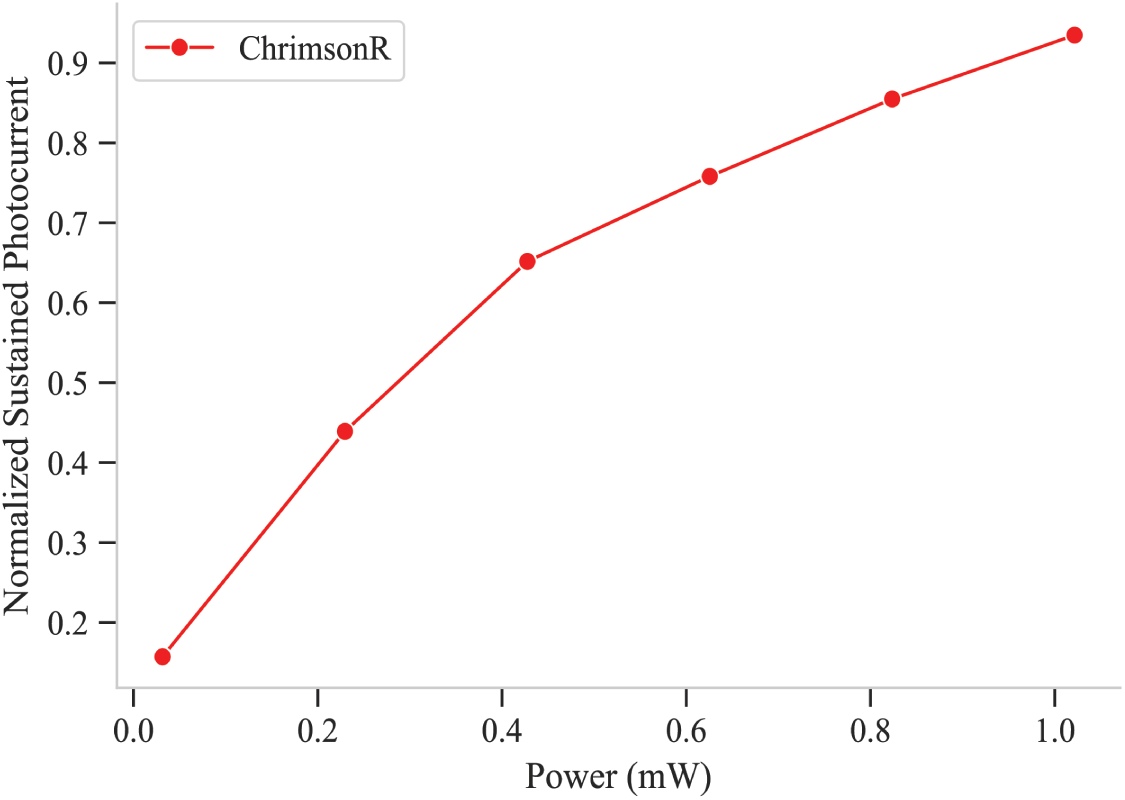
Intensity sweep of ChrimsonR (n=13).

**Figure A.7:**
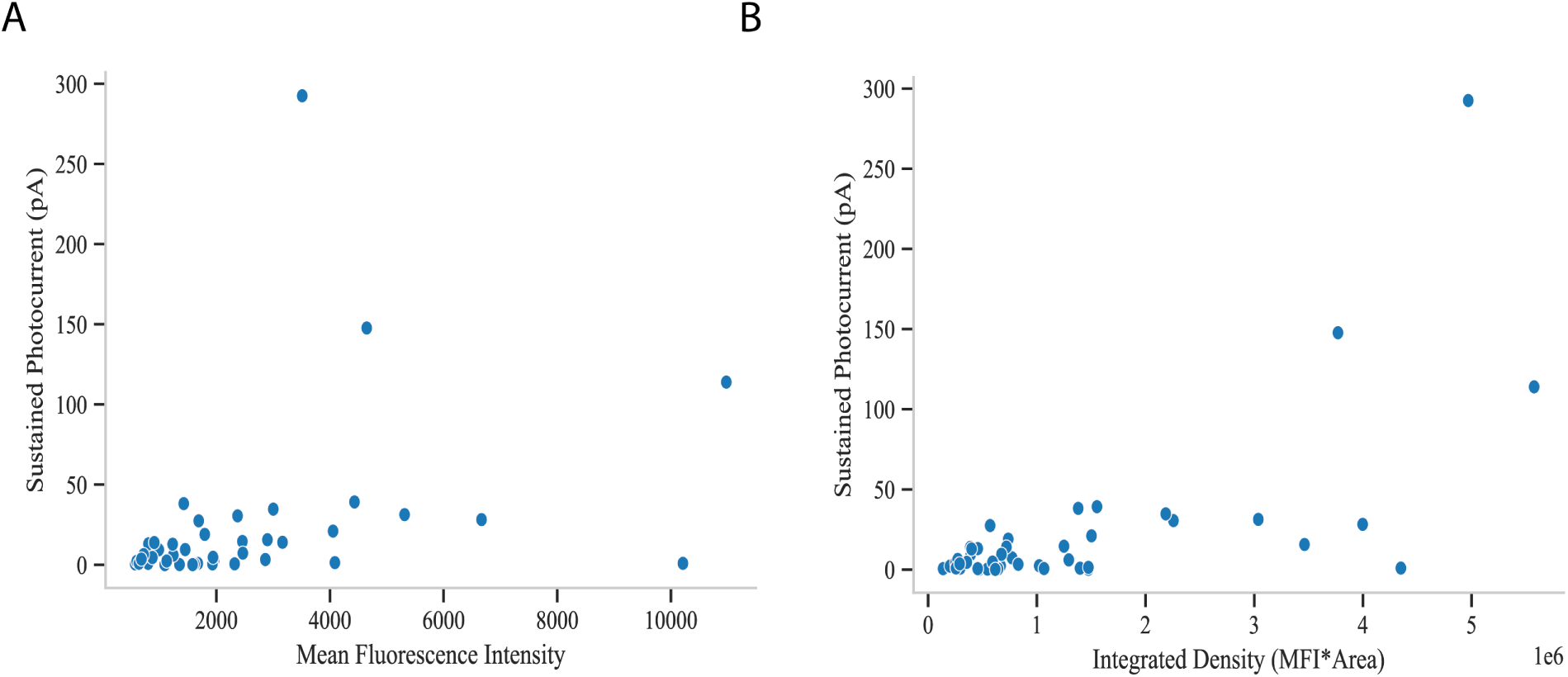
Sustained photocurrent as a function of fluorescent expression. (A) Mean fluorescence intensity versus sustained photocurrent (n=45). (B) Integrated density versus sustained photocurrent (n=45).

**Figure A.8:**
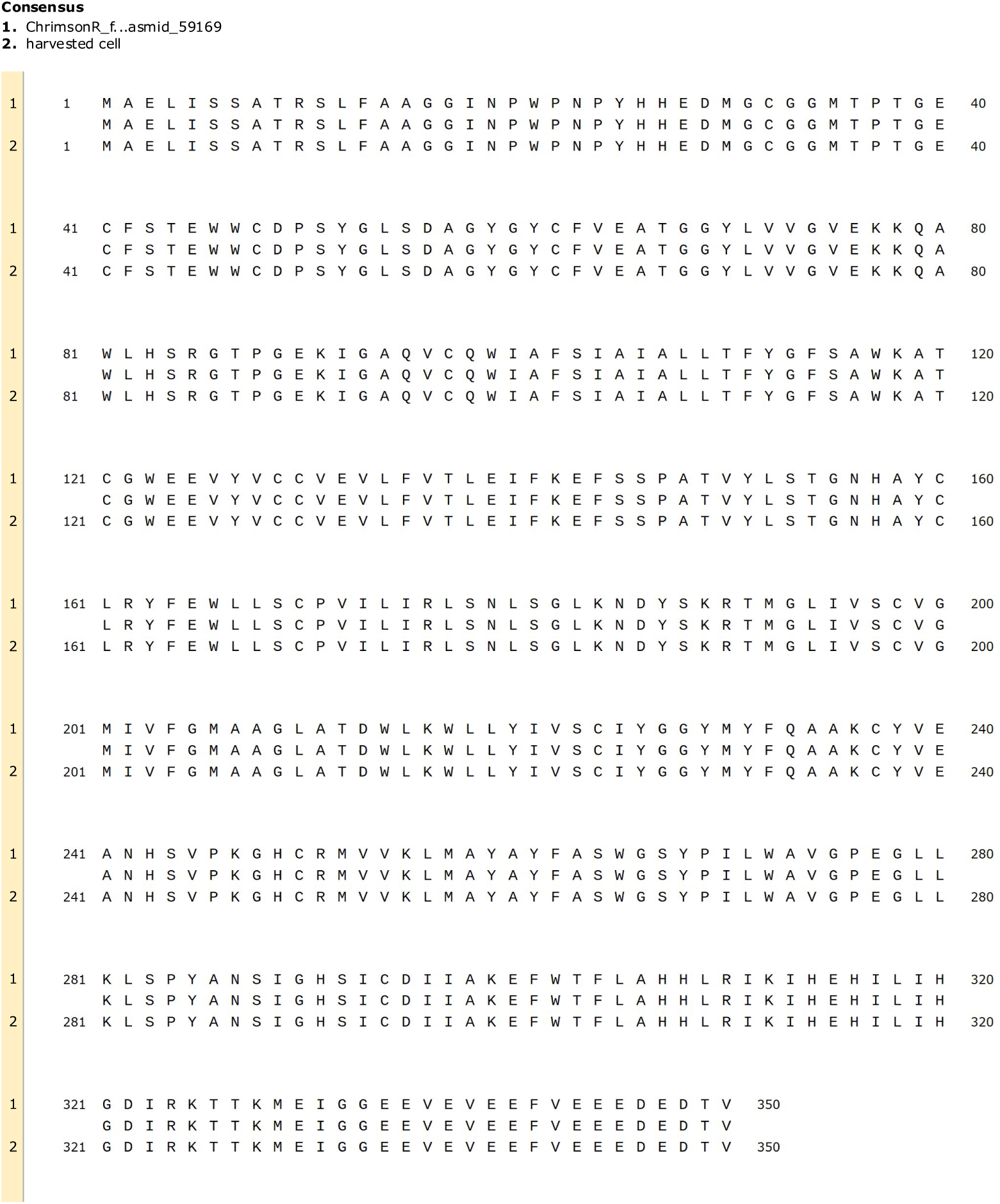
Example sequence alignment of harvested cell with known ChrimsonR sequence. Sequences were proven to be identical.

**Figure A.9:**
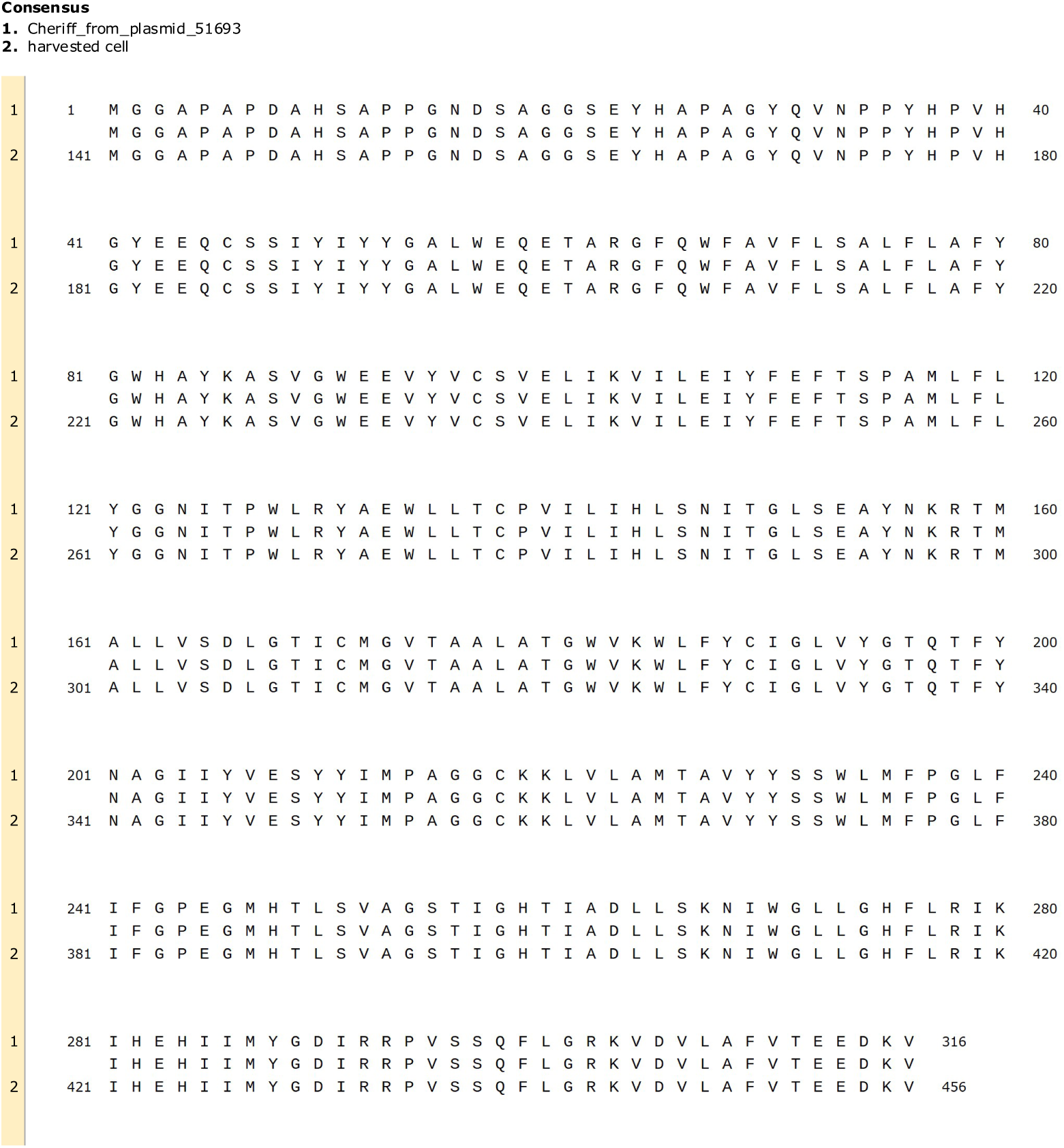
Example sequence alignment of harvested cell with known CheRiff sequence. Sequences were proven to be identical.

**Figure A.10:**
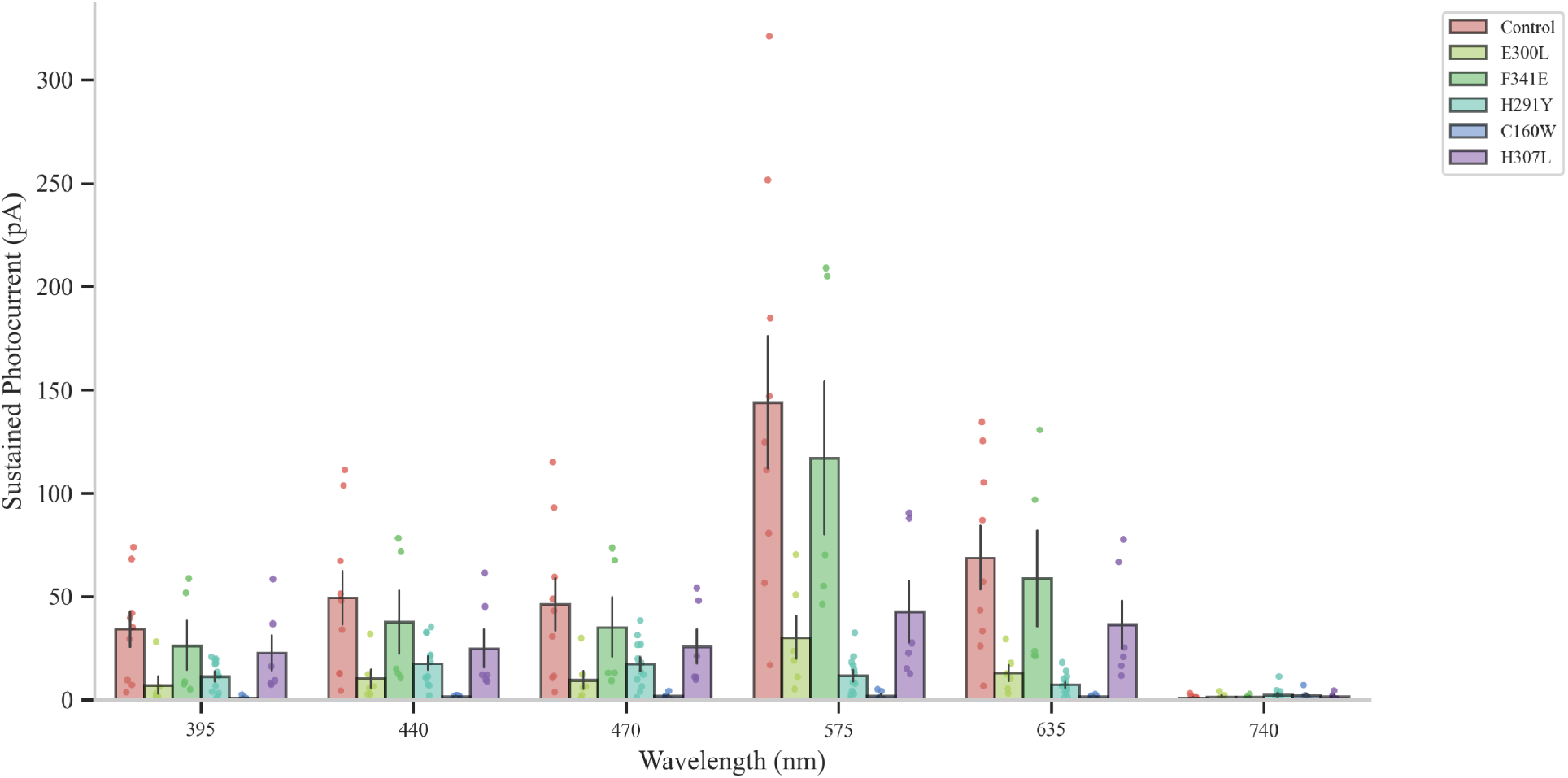
Baseline photocurrent spectra for ChrimsonR single substitution mutants. Sustained photocurrent amplitudes recorded for each variant across six wavelengths (395, 440, 470, 575, 635, and 740 nm). Each bar shows mean ± SEM; individual data points are overlaid.

**Figure A.11:**
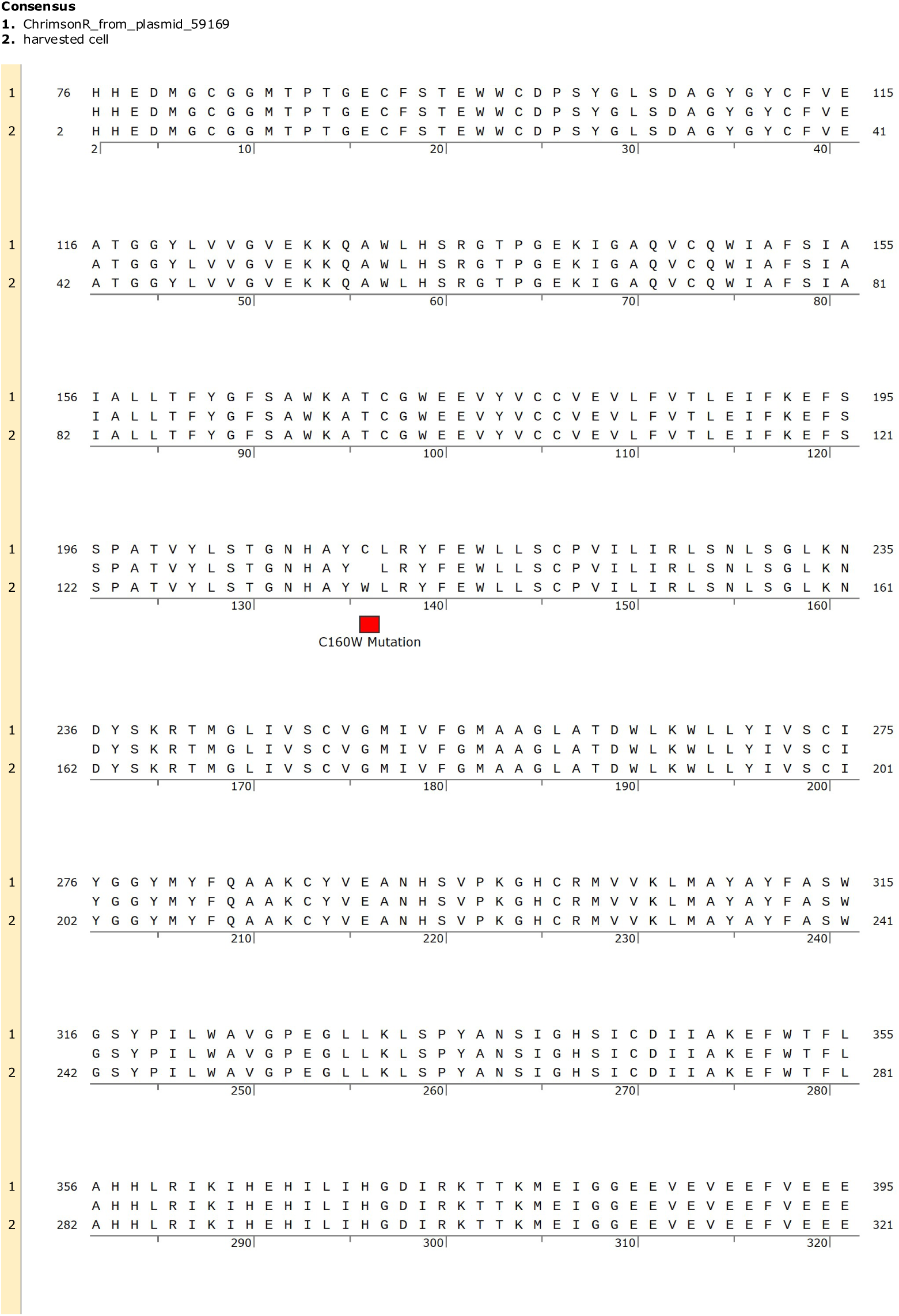
Example sequence alignment of harvested cell expressing C160W mutation with known ChrimsonR sequence. No unanticipated mutations were found.

**Figure A.12:**
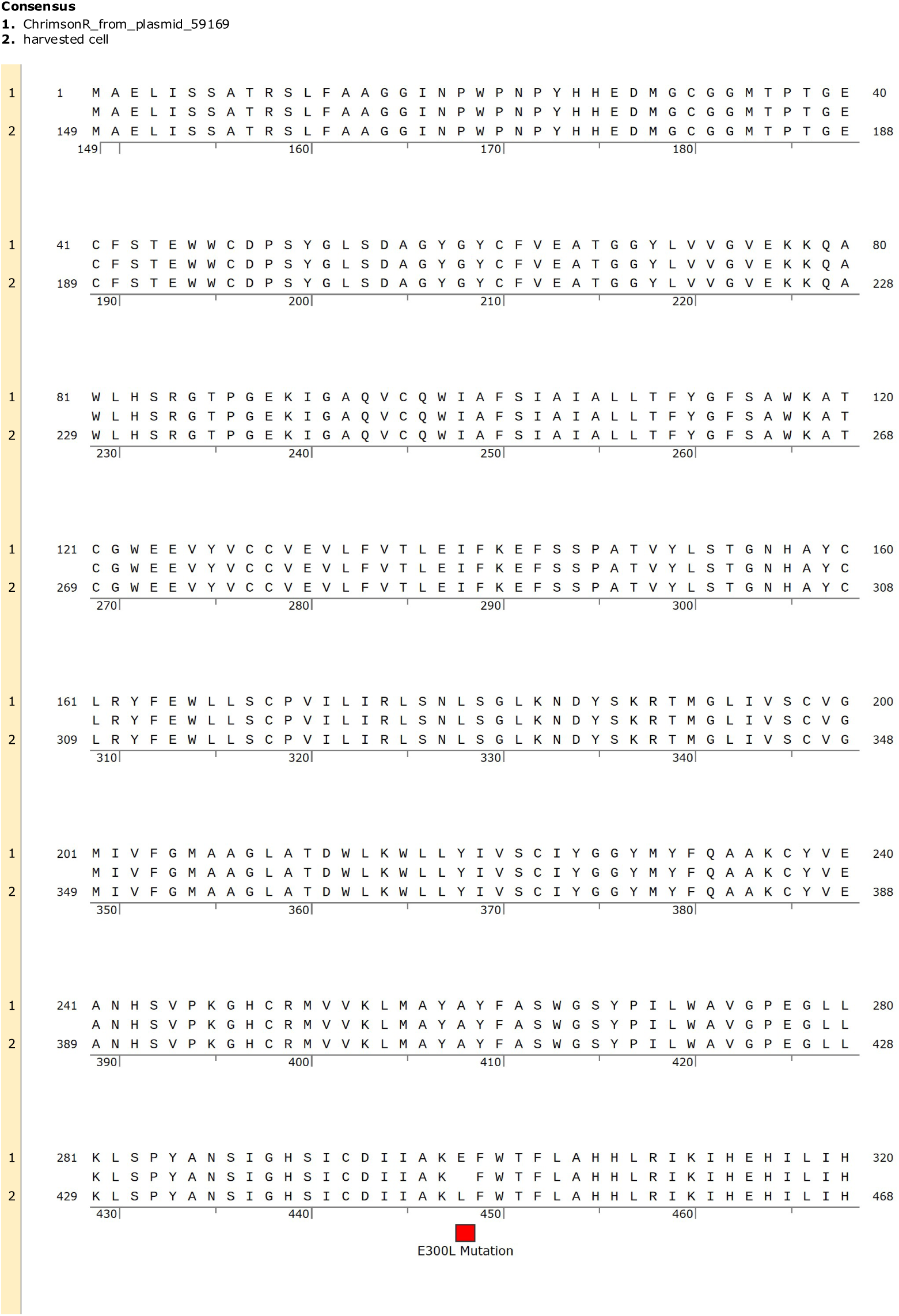
Example sequence alignment of harvested cell expressing E300L mutation with known ChrimsonR sequence. No unanticipated mutations were found.

